# Read counts from environmental DNA (eDNA) metabarcoding reflect fish abundance and biomass in drained ponds

**DOI:** 10.1101/2020.07.29.226845

**Authors:** C. Di Muri, L.L. Handley, C.W. Bean, J. Li, G. Peirson, G.S. Sellers, K. Walsh, H.V. Watson, I.J. Winfield, B. Hänfling

## Abstract

The sampling of environmental DNA (eDNA) coupled with cost-efficient and ever-advancing sequencing technology is propelling changes in biodiversity monitoring within aquatic ecosystems. Despite the growth of DNA metabarcoding approaches, the ability to quantify species biomass and abundance in natural systems remains a major challenge. Few studies have examined the association between eDNA metabarcoding data and biomass inferred by whole-organism sampling, mesocosms or mock communities, and the interpretation of sequencing reads as a measure of biomass or number of organisms is largely disputed.

Here we tested whether read counts from eDNA metabarcoding provide accurate quantitative estimates of fish abundance in holding ponds with known fish biomass and number of individuals.

eDNA samples were collected from two fishery ponds with high fish density and broad species diversity. In one pond, two different DNA capture strategies (on-site filtration with enclosed filters and three different preservation buffers versus lab filtration using open filters) were used to evaluate their performance in relation to fish community composition and biomass/abundance estimates. Fish species read counts were significantly correlated with both biomass and abundance, and this result, together with information on fish diversity, was repeatable when open or enclosed filters with different preservation buffers were used.

This research demonstrates that eDNA metabarcoding provides accurate qualitative and quantitative information on fish communities in small ponds, and results are consistent between different methods of DNA capture. This method flexibility will be beneficial for future eDNA-based fish monitoring and their integration into fisheries management.

## Introduction

Environmental DNA (eDNA) metabarcoding is revolutionising biomonitoring in aquatic environments (Harper et al., 2019; Jerde, 2019; Lawson Handley, 2015; Sigsgaard et al., 2019). This approach relies on the molecular identification of organisms whose genetic material has been collected, isolated and extracted from water. Species identification occurs after PCR with broad-range primers followed by High Throughput Sequencing and matching sequence reads against a reference database (see e.g. Valentini et al., 2016; Deiner et al., 2017 for an overview).

eDNA metabarcoding has been recently suggested as a complementary biomonitoring strategy for the European Union Water Framework Directive (WFD, 2000/60/EC) which requires member states to assess the ecological status of freshwater bodies. Currently established WFD methodologies include the morphological identification and counting of phytoplankton, phytobenthos and benthic invertebrates or gillnetting and electrofishing for fish (Hering et al., 2018). Yet traditional biomonitoring methods have limitations which may hamper species’ detectability or correct identification. They often lack broad applicability and they frequently impact on species’ welfare, such as the use of gillnets for fish (Radinger et al., 2019). eDNA metabarcoding has the advantage of detecting elusive and rare species, resolving cryptic species and identifying novel taxa through a non-invasive sampling approach (Blackman et al., 2017; Grey et al., 2018; Bylemans et al., 2019). The ease of eDNA collection also makes this approach suitable for remote location sampling, and the molecular identification of the genetic material does not require taxonomic expertise. eDNA metabarcoding has been shown to outperform established methods for the assessment of freshwater fish community composition (Civade et al., 2016; Hänfling et al., 2016; Valentini et al., 2016; Pont et al., 2018; Sard et al., 2019).

The ability of eDNA metabarcoding to provide information on abundance and biomass is more controversial, and uncertainties regarding the quantitative power of eDNA metabarcoding are still present among the scientific community and monitoring agencies (Fonseca, 2018; Lamb et al., 2019). This is particularly important given that species abundance is a crucial component of biodiversity surveillance and ecological monitoring schemes, and in view of ongoing biodiversity changes worldwide (Ficetola et al., 2018). Positive correlations between eDNA metabarcoding data (i.e. site occupancy or read counts) and fish abundance or biomass (as deduced by established surveys e.g. gill-netting) have been demonstrated in natural environments (Thomsen et al., 2012; Hänfling et al., 2016; Lawson Handley et al., 2019; Sard et al., 2019). However, estimates from established surveys also have their own biases and may not necessarily reflect true species abundance. Accurate data on organism-based measures of abundance from natural aquatic habitats are difficult to obtain without exhaustive sampling – such as draining down water bodies - and hence authentic comparisons with eDNA data in natural systems are, to our knowledge, still very rare.

A second key question is how replicable eDNA metabarcoding is with different field and laboratory protocols. Standardization of protocols may overcome this issue, but a “one-size fits all” protocol would be unrealistic (Ruppert et al., 2019). For instance, eDNA capture methods are often chosen based on factors such as proximity/accessibility of sampling locations and the availability of lab equipment. At present, enclosed filters are usually preferred for on-site processing, especially when remote locations are sampled, and storage buffers are used for DNA preservation within the encapsulated filter (Spens et al., 2017; Li et al., 2018; Takahashi et al., 2020). For field workers this approach would be logistically simple, less prone to contamination and much easier to integrate into monitoring programmes compared to laboratory based methods of eDNA capture. Open filter membranes allow a larger volume to be filtered, but suffer from field and transportation logistics, and are potentially more vulnerable to the risk of contamination (Li et al., 2018; Majaneva et al., 2018).

To evaluate the efficiency and suitability of different eDNA capture, a number of published studies have compared different approaches (precipitation versus filtration; on-site versus in laboratory), and a variety of filtration equipment, filters material and filters pore size (e.g. Deiner et al., 2015; Eichmiller et al., 2016; Lacoursière-Roussel et al., 2016; Minamoto et al., 2016; Djurhuus et al., 2017; Majaneva et al., 2018). Recent studies have also investigated the ability of different filter types (enclosed and open filters) and preservation methods (buffers and freezing) to provide quantitative estimates of eDNA using organisms’ biomass and abundance estimates from artificial stocked ponds (Li et al., 2018) or from in-field visual surveys (Takahashi et al., 2020). Evaluation of the quantitative performance of filter types and preservation methods based on absolute values of species biomass and numbers in natural environments would greatly contribute to the implementation of future eDNA-based surveys.

In the present study we tested whether eDNA metabarcoding can provide accurate information on the community composition and fish biomass and abundance in ponds that were drained as part of an invasive species eradication programme. During the drain down, all fish were counted, measured and weighed, providing absolute measures of species abundance and biomass, and so avoiding the biases of established techniques used in previous studies. Secondly, we tested whether estimation of fish abundance and biomass with eDNA metabarcoding is consistent between different methods of DNA capture, by comparing Sterivex enclosed filters preserved with three different buffers (ethanol, Longmire’s solution and RNAlater) and open filtration (using Mixed Cellulose Ester; MCE filters and a vacuum pump) followed by freezing preservation at -20°C.

## Methods

### Study site and collection of fish abundance and biomass data

The study was carried out at a UK fishing venue which consisted originally of three hydrologically-isolated stocked ponds (Upper, Middle and Lower Lake; Fig. 1A). This site was included in an Environment Agency (EA) eradication programme for non-native topmouth gudgeon (*Pseudorasbora parva*), as part of a wider government strategy to tackle invasive species in the UK (GB Non-Native Species Secretariat, www.nonnativespecies.org). In November 2016, during the eradication programme, a new pond of 0.2 ha (hereafter “New Lake”) was created, and the original three ponds drained. All fish over 150 mm total length from the original ponds were moved to the New Lake. During relocation, fish were individually checked for potentially hidden *P. parva* individuals in their gills and mouths. The original, empty ponds were partially refilled with water and treated with the piscicide rotenone by the EA to kill all potentially remaining specimens of *P. parva*. Original ponds were left fish-free for three months (from November 2016 to January 2017). On the 18^th^ January 2017, New Lake was completely drained and all fish were moved back to the original ponds. During fish re-allocation, individual fish were morphologically identified by experts, counted and weighed, hence the exact fish biomass and population size could be calculated for each species and water body. Following the fishery owner’s request, two of the original ponds (Upper and Lower Lake) became carp ponds, and they were re-stocked mainly with *Cyprinus carpio* and a few individuals of *Perca fluviatilis* and *Carassius carassius* x *C. carpio* hybrids. Middle Lake (0.3 ha) was re-stocked with 1,248 fish with a total biomass of 634,87 kg, equivalent to 2,116.23 kg/ha. The fish community included eight species and two hybrids with biomass and number of individuals ranging from 0.7 kg/1 individual (*Squalius cephalus*) to 240.6 kg/382 individuals (*Abramis brama*) (Fig. 1B; Table S1). New Lake fish community was then calculated as the sum of fish species and hybrids counted and weighed after fish re-allocation to the original ponds, and included a total number of twelve species and two hybrids with biomass and numbers ranging from 0.7-1 kg/1 individual (*S. cephalus* and *Acipenser spp*.) to 1,715.2 kg/483 individuals (*C. carpio*) (Fig. 1B; Table S1). Overall, New Lake contained 2,000 fish with a total biomass of 2,695.32 kg, equivalent to 13,476.6 kg/ha. Given the diverse fish communities of New Lake and Middle Lake, our eDNA metabarcoding analyses focused on these two ponds.

**Figure 1.**
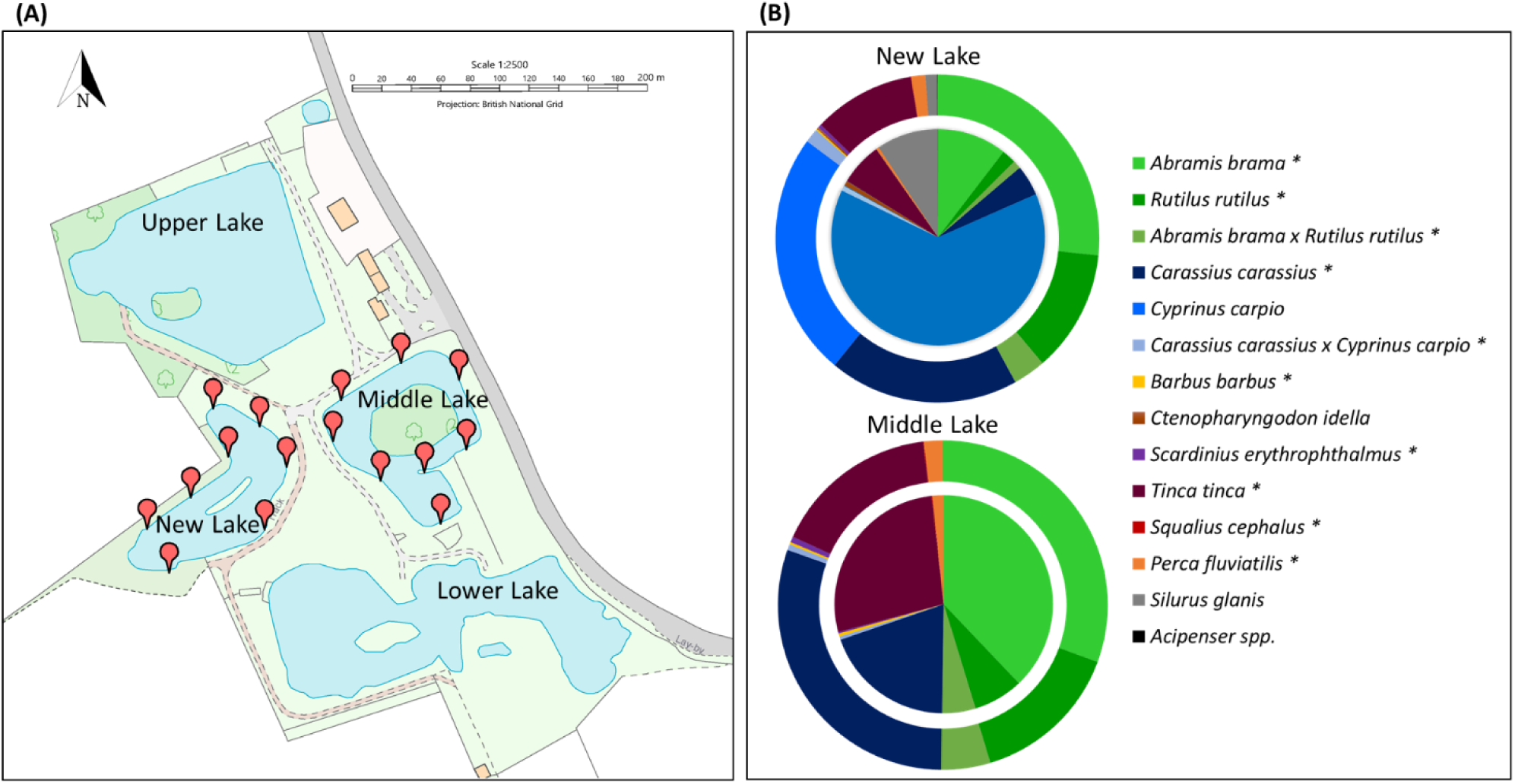
Map and fish diversity of the site surveyed. (A) Map of eDNA collection sites (in red) at the fishery venue. Map was downloaded and edited from Digimap (https://digimap.edina.ac.uk). (B) Fish species composition of the New Lake and Middle Lake after re-stocking (species with asterisk only). Ring pie charts (outer circles) show proportion of species composition by number of individuals; pie charts (inside circles) indicate proportion of species composition by fish biomass (kg).

### Water sample collection, filtration, and extraction

Water samples were taken on three separate occasions applying different strategies based on the goal of each occasion (see Fig. 2 for experimental design). New Lake was sampled the day before fish were transferred back to the original lakes (16^th^ of January 2017) using open filter membranes for eDNA capture (Fig. 2). We allowed one month after reintroductions for DNA dispersion in the water, and sampled Middle Lake on the 16^th^ and 17^th^ of February 2017, using replicated enclosed Sterivex filters and different preservation buffers (STX; Fig.2) and open filter membranes (MCE; Fig. 2).

**Figure 2.**
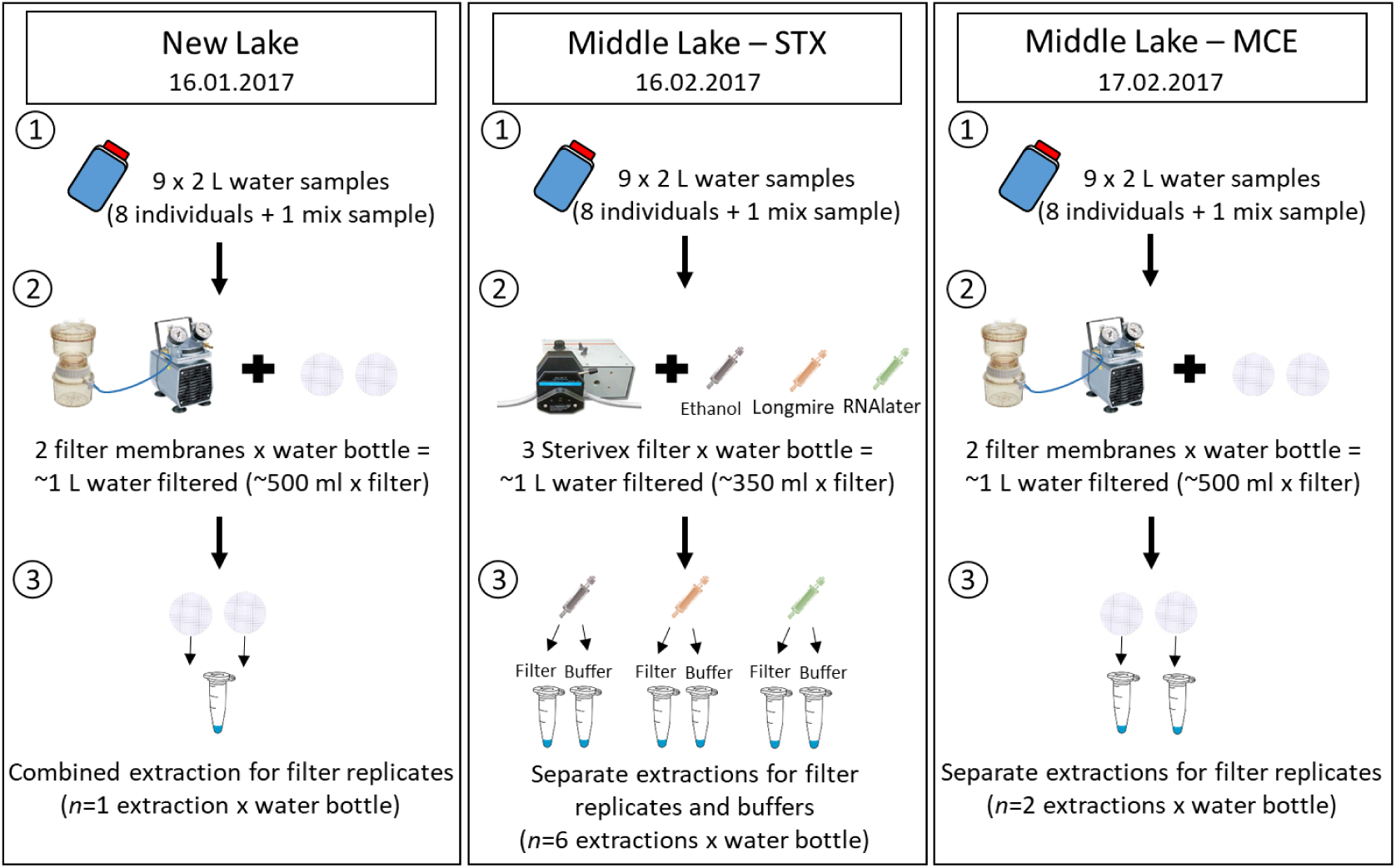
Experimental design. Panels show eDNA collection and processing strategies at different sampling occasions. Numbers within the panels indicate the workflow from water sampling (1) to filtration (2) and DNA extraction (3).

All precautions to avoid contamination were taken while sampling. Fieldwork equipment was sterilized using 10% v/v chlorine-based commercial bleach (Elliott Hygiene Ltd, UK) and sterile gloves (STARLAB, UK) were changed at each sampling location. Blanks, consisting of 2 L sampling bottles filled with ultra-purified water (Milli-Q), were included for each sampling occasion. Blanks were opened once in the field and then kept and processed alongside other water samples.

On each sampling occasion, eight 2 L water samples were collected equidistantly (∼30 m apart) around the perimeter of each pond (Fig. 1A). Samples were collected by hand at the water surface by pooling five 400 mL subsamples collected within a range of 5 m from the central location into a 2 L sterile plastic bottles (Gosselin™ Square HDPE, Fisher Scientific UK Ltd, UK). At each sampling occasion, immediately before filtration, a mixed sample was created using 200 ml aliquots from each of the eight water bottles collected in the field in order to evaluate differences of species detections with sampling strategies.

Samples for open filtration were placed inside cool boxes with ice packs, transported back to the laboratory and processed within 12 hours from collection. eDNA was captured onto 0.45 μm mixed cellulose ester membranes (MCE, 47 mm diameter, Whatman, GE Healthcare) using a vacuum-pump and Nalgene™ filtration units. Filtration was stopped after 45 min and approximately 500 ml of water was filtered through each of two MCE open filter membranes per sample (i.e. 1 L of the 2 L total sample was filtered). Filter membranes were then stored in sterile 50 mm Petri dishes (Fisher Scientific UK Ltd, UK) sealed with parafilm (Bemis™, Fisher Scientific UK Ltd, UK) and kept at -20**°**C until DNA extraction.

Sterivex filtration was carried out in the field. eDNA was captured using 0.45 μm Sterivex filter units (PVDF membrane, Merck Millipore) connected to a peristaltic pump (Easy Load II Peristaltic Pump, In-situ Europe Ltd, UK). On-site filtration was also carried out until an individual filter became clogged, otherwise it was stopped after 45 min. Approximately 350 ml were filtered through each Sterivex filter and three Sterivex units were used per sample. Each filter was then preserved using 2 ml of one of three different buffers: ethanol (≥ 99.5% v/v), Longmire’s solution, and RNAlater.

All DNA extractions were carried out using the Mu-DNA protocol for water samples following adaptation for Sterivex as recommended in Sellers et al. (2018), and the DNA was eluted into 100 μl of TE buffer. Filter replicates of open membranes from New Lake were co-extracted by placing both filters in a single tube for bead milling, whereas, to compare metabarcoding results of open membranes from the Middle Lake-MCE sampling, filter replicates were extracted separately (Fig. 2). For Sterivex units, DNA from buffers and filters was extracted separately as previous studies have shown that DNA can become suspended in the buffer (Spens et al., 2017; Fig. 2). After extractions, nucleic acid yield and purity were checked on a Nanodrop 1000 spectrophotometer (Thermo Fisher Scientific).

### Library preparation

Library preparation included a two-step PCR with a nested-tagging approach as described in Li et al. (2019a, 2019b). In the first round of PCR, indexed primers targeting a 106 bp region within the 12S fragment were used (Riaz et al., 2011; Kelly et al., 2014). The first round of PCRs was performed in a final reaction volume of 25 μl including 12.5 μl of Q5^®^ Hot-Start High-Fidelity 2X Master Mix (New England Biolabs^**®**^ Inc., MA, USA), 1.5 µl of each indexed primer (10 µM; Integrated DNA Technologies, Belgium), 7.5 µl of molecular grade water (MGW; Fisher Scientific UK Ltd, UK) and 2 μl of DNA template at the original sample concentration. In order to avoid cross-contamination between samples as a consequence of evaporation and/or aerosols, reactions were prepared in 8-strip tubes with individually attached caps and covered with a drop of mineral oil (Sigma-Aldrich Company Ltd, UK). Amplifications were performed on Applied Biosystems^**®**^ Veriti thermal cyclers (Life Technologies, CA, USA) with the following conditions: initial denaturation at 98°C for 5 min; 35 cycles of 98°C for 10 sec, 58°C for 20 sec and 72°C for 30 sec; final elongation step at 72°C for 7 min. Eighty-one samples, eight collection blanks, six PCR negatives (Molecular Grade Water, MGW), and four positives (genomic DNA [0.05 ng/ μl] from cichlid species not occurring in the UK, *Astotilapia calliptera* and *Maylandia zebra*) were amplified in triplicate. Amplicons were checked on 2% agarose gels stained with 10,000X GelRed Nucleic Acid Gel Stain (Cambridge Bioscience, UK). Gels were imaged using Image Lab Software (Bio-Rad Laboratories Ltd, UK) to visually check for contamination in blanks/negatives, presence of target band and consistency of results among PCR replicates.

After visualization, PCR triplicates were combined and samples belonging to the same collection site were pooled and normalized using different volumes as deduced from strength of PCR products on gels (no/very faint band = 10 µl, faint band = 7.5 µl, bright band = 5 µl) using 1 μl of the positive samples and 5 μl of blanks/negatives for each pool (Alberdi et al., 2018).

Amplicon pools were cleaned using a double-size selection magnetic beads protocol (Bronner et al., 2013) with a ratio of 0.9X and 0.15X of magnetic beads (Mag-Bind^®^ RXNPure Plus, Omega Bio-tek Inc, GA, USA) to PCR products. Bead purification was followed by a second amplification where Illumina tags were added to each pool. Second PCRs were run in duplicate in a final reaction volume of 50 µl using 25 µl of Q5^®^ Hot-Start High-Fidelity 2X Master Mix (New England Biolabs^**®**^ Inc., MA, USA), 3 µl of each Illumina tag (10 µM; Integrated DNA Technologies, Belgium), 15 µl of MGW (Fisher Scientific UK Ltd, UK) and 4 µl of pooled templates. PCRs consisted of: 95°C for 3 min; 8 cycles of 98°C for 20 sec and 72°C for 1 min; and 72°C for 5 min. PCR products were checked on a 2% agarose gel alongside their non-tagged products to check for size differences after tag addition. A second double-size selection bead purification was carried out with a ratio of 0.7X and 0.15X of magnetic beads/PCR products. Tagged amplicon pools were quantified using the Qubit™ 3.0 fluorometer and a Qubit™ dsDNA HS Assay Kit (Invitrogen, UK) and pooled with equimolar concentrations into a unique library. The final library was checked for size and integrity using the Agilent 2200 TapeStation and High Sensitivity D1000 ScreenTape (Agilent Technologies, CA, USA) and quantified using qPCR with the NEBNext^**®**^ Library Quant Kit for Illumina^**®**^ (New England Biolabs^**®**^ Inc., MA, USA). Following qPCR, 13 pM library was loaded on the Illumina MiSeq^®^ with 10% PhiX using a 2 × 300 bp V3 chemistry (Illumina Inc., CA, USA).

### Bioinformatics and statistical analyses

Raw sequencing data were demultiplexed using a custom Python script and subsequently analyzed with metaBEAT (metaBarcoding and Environmental Analysis Tool) v0.97.11 (https://github.com/HullUni-bioinformatics/metaBEAT), an in-house developed pipeline. Quality trimming, merging, chimera detection, clustering and taxonomic assignment against a custom-curated 12S reference database (Hänfling et al., 2016) containing sequences for all UK freshwater fish species were performed. The number of reads assigned to species (i.e. read counts) was used for downstream analyses in R v.3.5.1. (R Core Team 2018).

Total read count per sample was calculated as the sum of assigned and unassigned reads. The proportion of reads assigned to each fish species over the total read counts was then calculated on a sample by sample basis. A low-frequency noise threshold of 0.001 (0.1%) was applied across the dataset to reduce the probability of false positives arising from cross-contamination or tag-jumping (De Barba et al., 2014; Hänfling et al., 2016). Based on the level of contamination found in sampling/filtration blanks and PCR negatives, a second arbitrary threshold was applied and all records occurring with less than 50 reads assigned were removed.

Morphological identification of fish species revealed that a substantial amount of F1 hybrids (Fig. 1; *C. carassius* x *C. carpio* and *A. brama* x *Rutilus rutilus*) were present. As community eDNA approaches are unable to differentiate hybrids from parental species these were grouped together for the purpose of our correlation analyses; i.e. data on biomass/abundance and eDNA read counts/site occupancy for hybrids and their parental species were pooled.

As read counts and site occupancy data were not normally distributed, Spearman’s rank correlation coefficient was used to calculate correlations between biomass/abundance data and species average read counts and site occupancy for filter types and treatments. Graphs were plotted using *ggplot2* (Wickham, 2016) and correlation coefficients and significant levels displayed using functions in *ggpubr* (Kassambara, 2018). Species site occupancy was calculated as the number of filter replicates with positive detections over the total number of filter replicates collected and processed using the same treatment (*n*=8).

VEGAN package v2.5-4 (Oksanen et al., 2019) was then used to test differences of fish communities between filter types (Sterivex and open membranes) and treatments (preservation buffers and freezing). *Betadisper* was used to investigate compositional variance of each group, and homogeneity of group dispersions was tested using ANOVA. Distances from the centroids of each treatment and the variance within treatment were visualized with a Principal Coordinates Analysis (PCoA). To test groups for compositional differences, a permutational multivariate analysis of variance (PERMANOVA), with replicates nested into each filter type, was carried out using the *adonis* function. Tests were performed on a square-root transformed abundance-weighed dissimilarity matrix (Bray-Curtis) of species composition.

Lastly, sample-based species accumulation curves (SACs) were built using the function *specaccum* for each filter type and replicate.

Details of protocols, bioinformatics, R script and supplementary material used for the analyses can be found at: DOI 10.17605/OSF.IO/ZWPSQ.

## Results

### Sequencing outputs and bioinformatics

The total number of paired-end sequences across 98 samples (81 eDNA samples and 17 controls) was 10,751,170. Of these, 6,398,530 sequences passed the trimming quality filter and 92% were subsequently merged. 3,389,668 sequences remained after chimera detection and clustering. Excluding the cichlid species used as positive controls, 16 Operational Taxonomic Units (OTUs), and 1,314,623 sequences were identified as fish taxa, with 100% match to the custom-curated fish reference database with thirteen OTUs remaining after applying the thresholds. All fish OTUs were identified to species level with the exceptions of records matching the family Percidae. Percidae records were manually assigned to *P. fluviatilis* as this was the only species of the family identified in the study area by whole-organism sampling.

*P. parva* reads found in two Middle Lake-STX samples (279 and 148 reads) were also excluded from further analyses as after eradication this species was not physically present at the site surveyed.

### eDNA metabarcoding fish diversity

OTUs from eleven of the twelve fish species translocated to New Lake were detected in eDNA samples, but two records were removed after applying thresholds. Sequences from the following taxa were detected at all eight sites within New Lake: *A. brama, C. carassius, C. carpio, P. fluviatilis, R. rutilus, Silurus glanis* and *Tinca tinca* (Fig. 4, S1) with *C. carpio* showing the highest read counts (about 40,000) and other species reads ranging from 1,831 of *S. glanis* to 23,618 of *A. brama* (Fig. 3, S1). In addition, *Barbus barbus* was detected at two sites (202 reads), and *Ctenopharyngodon idella* at one site (71 reads) (Fig. 3, 4, S1). The presence of *Scardinius erythrophthalmus* was found at two sites with a low number of reads (38 and 25 reads) and, therefore, removed after applying the filter threshold (see metaBEAT raw data, Table S2). Taxonomic assignment based on our reference database failed to detect *Acipenser spp*., yet 79 reads (at one site) matched the family Acipenseridae during the unassigned blast against GenBank, however this record was excluded from further analyses (see unassigned blast data, Table S3).

**Figure 3.**
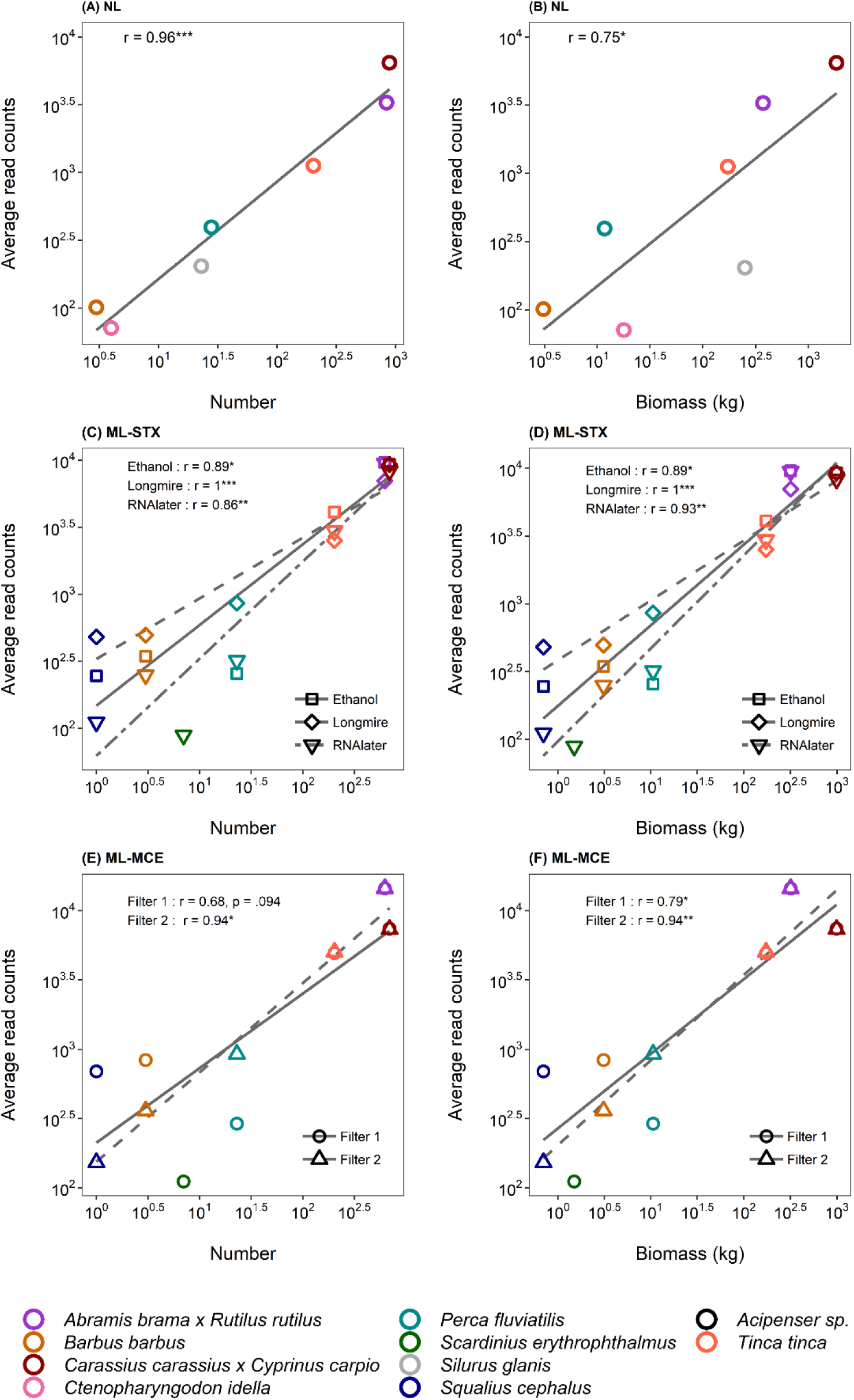
Correlations between eDNA metabarcoding read counts and fish abundance/biomass. Scatterplots showing Spearman’s correlations of fish species average read counts with abundance (number of individuals, on the left) and biomass (kg; on the right) at different sampling occasions. Panel (A) and (B) Spearman’s correlations for New Lake; (C) and (D) Spearman’s correlations for Middle Lake with Sterivex filters; (E) and (F) Spearman’s correlations for Middle Lake with open filter membranes (MCE). Plot axes were log transformed for better visualization. Significant codes: ***0.001; **0.01; *0.05.

**Figure 4.**
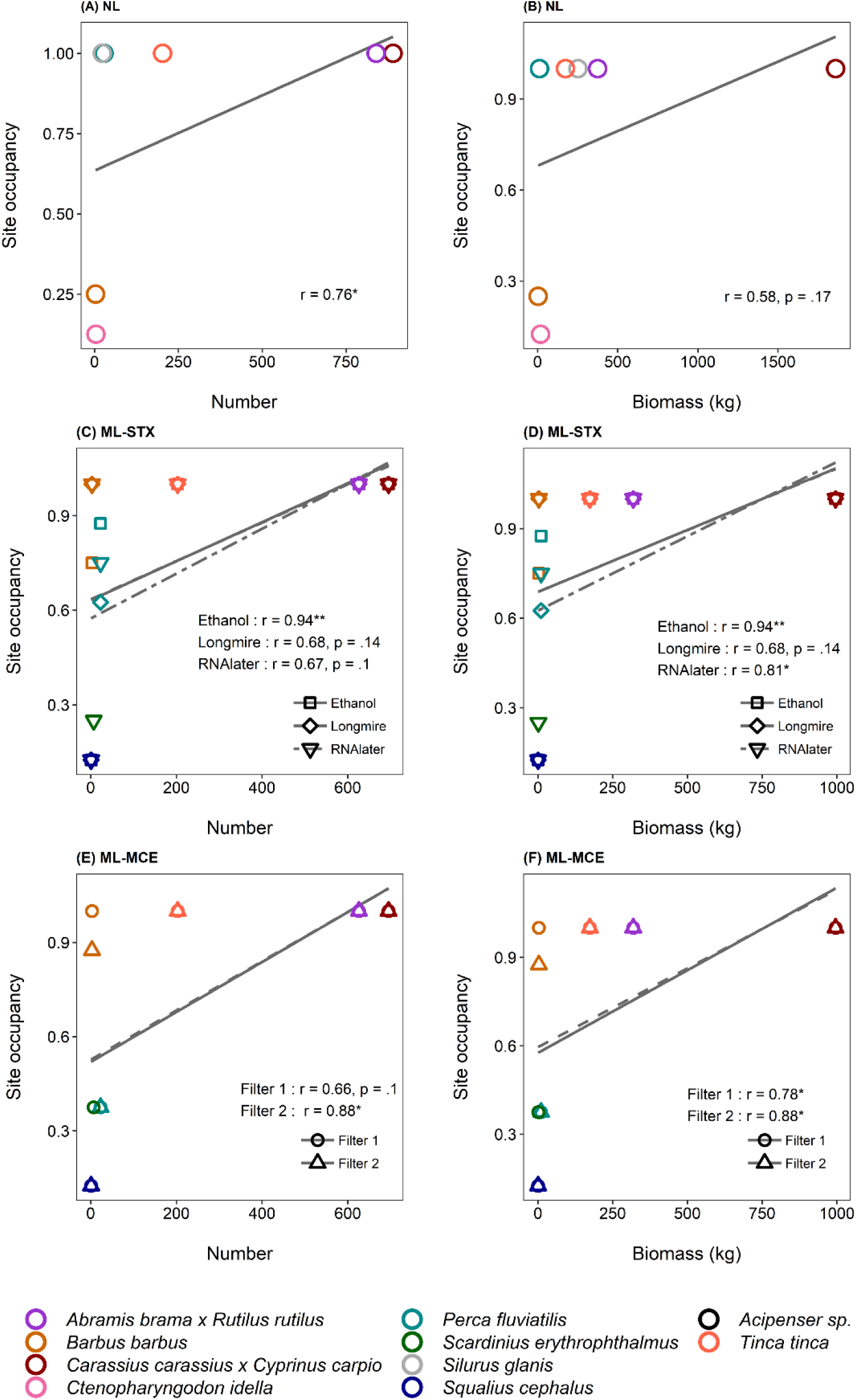
Correlations between eDNA metabarcoding site occupancy and fish abundance/biomass. Scatterplots showing Spearman’s correlations of fish species site occupancy with abundance (number of individuals, on the left) and biomass (kg; on the right) at different sampling occasions. Panel (A) and (B) Spearman’s correlations for New Lake; (C) and (D) Spearman’s correlations for Middle Lake with Sterivex filters; (E) and (F) Spearman’s correlations for Middle Lake with open filter membranes (MCE). Significant codes: ***0.001; **0.01; *0.05. Note: mix samples were not included in the analyses.

All nine possible OTUs corresponding to the species reintroduced were detected beyond threshold limits in Middle Lake in both sampling occasions (16^th^ and 17^th^ of February). Eight OTUs (*A. brama, R. rutilus, C. carassius, C. carpio, T. tinca, B. barbus, P. fluviatilis, S. cephalus)* were detected in both Middle Lake-STX and Middle Lake-MCE, and with all filter replicates (Fig.3, 4, S1). Five of these fish OTUs (*A. brama, R. rutilus, C. carassius, C. carpio, T. tinca*) showed high site occupancy (all sites occupied) and number of reads assigned (Fig. 3, 4, S1). Detection was less consistent for one of the two least abundant species, *S. erythrophthalmus*. In Middle Lake-STX, *S. erythrophthalmus* was only detected in one filter replicates preserved with RNAlater (266 reads), and in Middle Lake-MCE, in one filter membrane replicate 1 (333 reads; Fig. 3, 4, S1).

### Correlation between eDNA and biomass/abundance data

We evaluated the relationship between fish eDNA read counts/site occupancy of different filter replicates and fish biomass and abundance in New Lake and Middle Lake.

We observed significant positive correlations between fish read counts and fish biomass (r = 0.75; p = 0.052; Fig. 3B) and between read counts and number of individuals (r = 0.96; p < 0.001; Fig. 3A) for samples collected from New Lake.

Spearman’s correlations were calculated separately for each filter type (Sterivex/filter membranes) and filter replicate for samples collected from Middle Lake (ML-STX, ML-MCE). Fish read counts for all replicates and filters were positively correlated to both fish biomass and number of individuals. The highest associations were observed when read counts of Sterivex filter replicates were compared with biomass (Ethanol: r = 0.89, p = 0.019; Longmire: r = 1, p < 0.001; RNAlater: r = 0.93, p = 0.0025; Fig. 3D), and number of individuals (Ethanol: r = 0.89, p = 0.019; Longmire: r = 1, p < 0.001; RNAlater: r = 0.86, p = 0.014; Fig. 3C).

For open filter membranes (ML-MCE), there was a significant correlation between read counts and biomass for both filter replicates (r = 0.79, p = 0.036; r = 0.94, p = 0.0048; Fig. 3F) and between read count and number of individuals for filter 2 (r = 0.94, p = 0.048; Fig. 3E), but the correlation between read count and number was not significant for filter 1 (r = 0.68, p = 0.094; Fig. 3E).

Positive but weaker correlations of New Lake eDNA samples were observed when species site occupancy was associated with fish biomass (r = 0.58, p = 0.17; Fig. 4B) and number of individuals (r = 0.76, p = 0.049; Fig. 4A).

Fish site occupancy of Middle Lake filter replicates (ML-STX, ML-MCE) was also positively correlated to both fish biomass and number of individuals with, however, weaker associations. Correlation coefficients and significance of the Spearman’s correlations varied between filter replicates of both filter types. The strongest associations were observed when site occupancy of Sterivex filters preserved with ethanol were correlated with number of individuals and biomass (Ethanol: r = 0.94, p = 0.0051; Fig. 4C, 4D), but also when site occupancy of open filter membrane replicate 2 were associated with fish species biomass and fish number (r = 0.88; p = 0.021; Fig. 4E, 4F).

### Effect of sampling and filtration strategies on fish community eDNA data

To evaluate the effect of different sampling strategies the mean species richness of individual samples was compared to the species richness of the mixed sample at each sampling occasion and treatment (Fig. 5A). Overall, the number of fish species detected in the mixed samples was very close, and most of the time higher, than the average number of species detected in individual field samples with the only exception of MCE filter replicate 2 (Fig. 5A). The fish species not represented in the mixed samples were often the low-occurrence taxa of the sites surveyed, and generally, excluding MCE filter 2, a number of two fish species were missing in the mixed samples. For example, in the New Lake mixed sample *B. barbus* and *C. idella* were not detected. *S. cephalus* and S. e*rythrophthalmus* were not represented in Middle Lake-STX (ethanol, RNAlater and Longmire’s preservation) nor in Middle Lake-MCE filter 1 and 2 with the latter one additionally missing *B. barbus* and *P. fluviatilis*.

**Figure 5.**
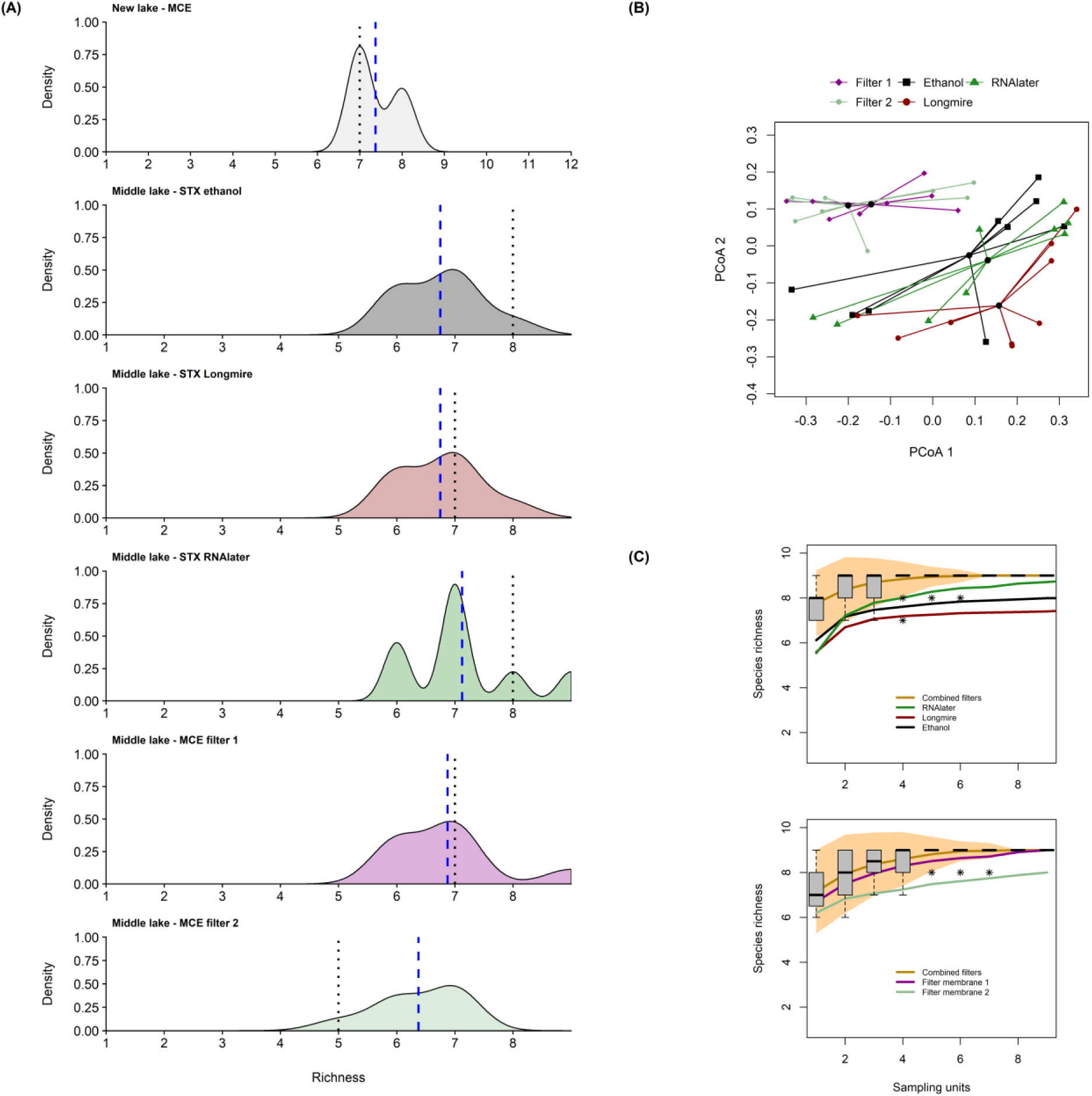
eDNA metabarcoding fish community plots for different filter types and treatments. (A) Density plots of species richness at different ponds and filtration strategies. The dashed blue line shows the species richness mean of eDNA samples (*n*=8), the dotted black line shows species richness of pooled samples collected at each sampling occasion. (B) PCoA plot showing distances from centroids of filter types (MCE and Sterivex) and treatments (buffers and freezing). Distances from centroids were calculated upon a dissimilarity matrix (Bray-Curtis) of fish species read counts. (C) Species accumulation curves for filter replicates of enclosed filters preserved with buffers (top) and open filter membranes with freezing preservation (bottom). In both figures, gold curves representing “Combined filters”, are calculated based on the sum of species when filter replicates of the same samples are combined. 95% confidence intervals refer to the “Combined filters” curves and boxplots of the same curves show distribution of species diversity as inferred from the method “random”, which add sites in random order and was used for the SACs.

Species accumulation curves of both Sterivex and MCE filters showed that approximately six samples are required to detect all fish species when filter replicates are combined (Fig. 5C). SACs of single filter replicates for Sterivex filters showed higher rates of species detection with RNAlater preservation compared to Longmire’s or ethanol preservation (Fig. 5C). For the open filters, most of the species were recovered with the first filter membrane, with only a slight improvement in detection rate when the second membrane was included (Fig. 5C).

There were no differences between fish community composition of different filter types (ANOVA *F* = 0.8521, *p* = 0.3611; Fig. 5B) or filter replicates (ANOVA *F* = 0.6495, *p* = 0.6305; Fig. 5B).

There was no significant difference between centroids of Middle Lake fish communities described by eDNA metabarcoding when using different filter types (PERMANOVA; R^2^ = 0.23278; p = 0.7231) or different preservation methods (buffers and freezing; R^2^ = 0.03795; p = 0.7231). However, more of the variation (approximately 23%) was explained by the use of different DNA capture methods (MCE versus Sterivex), compared to within filter treatment (∼3.8%).

## Discussion

With the advent of the next-generation eDNA-based monitoring surveys there is a growing interest in whether eDNA metabarcoding can generate accurate semi-quantitative data. Previous studies in natural environments have focussed on indirect estimates of fish abundance from established surveys which have their own inherent biases. Here, we used absolute data on fish abundance and biomass from drained ponds and found that read counts from eDNA metabarcoding consistently correlate with both fish numbers and biomass. Moreover, the present study suggests that the use of different eDNA capture (Sterivex vs. MCE open filters) and storage methods (buffers and freezing) produce repeatable results of fish diversity, composition and biomass/abundance estimates. We additionally show that the collection of spatial and filter replicates enhances species detection probability for rare species, thus sample coverage and replication are an important consideration in experimental design.

### Fish species detection

All 12 fish taxa were successfully detected in both fishery ponds surveyed with the only exception of *S. cephalus* in New Lake (single specimen of 0.7 kg; Fig. 1B, Table S1). Our findings are in line with other studies corroborating the ability of eDNA metabarcoding to describe fish diversity in lentic environments (Civade et al., 2016; Hänfling et al., 2016; Li et al., 2018; Lawson Handley et al., 2019; Zhang et al., 2020). The appropriate sampling effort, such as volume of water and spatial replicates collected, may vary according to the waterbody features (i.e. surface area, heterogeneity, species richness) and other environmental and biological factors (Civade et al., 2016; de Souza et al., 2016; Lawson Handley et al., 2019). In this study, the collection of eight, evenly distributed 2 L water samples from the ponds shore provided sufficient coverage of the fish community of the ponds surveyed. Eight 2 L samples collected from the edge of Middle Lake were appropriate for species detection at both sampling occasions and for the different filter types used. In fact, rarefaction curves (Fig. 5C) demonstrated that when filter replicates are combined, six 2 L water samples are sufficient to unveil the total fish composition of this intensively stocked and small size pond (0.3 ha and 2,116.23 kg/ha of fish density; Fig. 5A, C). In line with other eDNA metabarcoding studies, we suggest that nearshore sampling provides adequate species coverage as previously observed in larger and deeper lentic environments with complex fish species assemblages where a greater number of species has been detected inshore as opposed to offshore waters (Hänfling et al., 2016; Zhang et al., 2020). Here, we additionally highlight that an adequate sampling effort is paramount for describing species occurrence within a water body. In small, shallow lentic systems eDNA is thought to be homogeneously distributed in the water (Thomsen et al., 2012; Evans et al., 2017) even though the signal strength may increase closer to its source (Li et al., 2019b). Yet, we demonstrated that eDNA concentration of low-abundant species DNA is very localized, hence intensive sampling efforts and the collection of an adequate number of replicates is required to detect low-occurrence taxa. For example, our mixed samples (pooled water aliquots of field samples) consistently detected the common fish species at all sampling occasions, but failed to detect individuals or low-abundant taxa. Spatial pooling is therefore inefficient for detailed biodiversity surveillance as suggested by Zhang et al. (2020), who, on a larger spatial scale with higher number of PCR replicates, still found reduced OTUs detection in mixed water samples. In line with these results, we demonstrated that eDNA detection rate is enhanced with spatial and technical replication as well as with the increased water volume filtered.

Of particular interest is the detection of *P. parva* DNA in Middle Lake samples as this invasive species was the target of the eradication programme and present in extremely high abundance before the ponds were drained and treated with piscicide. The persistence of *P. parva* as living organisms within the pond appears extremely unlikely due to the effective eradication methods used in combination with the relatively small size of the water body (Britton et al., 2008; Genovesi and Carnevali, 2011). Furthermore, *P. parva* has not been recorded in these lakes since the eradication programme. Contamination could have occurred during the water sampling or in the laboratory resulting in false positive detection (e.g. Hänfling et al. 2016). However, no *P. parva* DNA was found in any of the control samples or in the water samples from New Lake. A possible explanation for this record is that *P. parva* eDNA originated from sediment resuspension in the water column during sampling or from carcasses remaining at the bottom of the pond. eDNA is known to be less concentrated and less persistent in water compared to sediment, where it remains detectable for over three months also when species are removed from the system (Turner et al., 2015). A further reasonable option would be to consider this result as a true record even if we have no evidence that the species re-colonised the pond after the eradication. Previous studies have suggested that *P. parva* may suffer from recruitment failure and local extirpation when population numbers are low due to human or natural disturbance (Coop et al., 2007; Davidson et al., 2017). Therefore, when monitoring the success of eradication attempts, extreme caution should be taken with false-positive or false-negative detections for the target species and the use of conventional methods to corroborate eDNA detections has been recommended (Davison et al., 2019; Robinson et al., 2019).

### Read counts correlate to biomass and abundance

To our knowledge, this is the first published study to date where the correlation between eDNA metabarcoding data and actual measures of species biomass and abundance in semi-natural lentic systems has been investigated. Our eDNA metabarcoding results accurately reflect relative abundance patterns and reveal positive and strong correlations between read counts and fish species biomass (weight) and number of individuals (Fig. 3). Recently Kelly et al. (2019) demonstrated that when amplification efficiency is high in PCR-based studies, proportional indices of eDNA reads capture trends in taxon biomass with high accuracy. Our study supports these findings as we found that the species read counts was an accurate quantitative tool to describe taxon biomass and abundance from taxon-specific read-abundance.

Positive associations were observed between species site occupancy and fish biomass/numbers, however, less significant than correlations with read counts (Fig. 4). In our study system, the relatively small size of the water bodies surveyed, coupled with the high fish densities, resulted in relatively homogeneous distribution of the common species’ eDNA, generating a better representation of fish biomass and abundance when read counts were used for quantitative inferences. In larger and heterogeneous lentic environments, the spatial variation of the species’ eDNA signal is likely to be as or more reliable than read counts for quantitative estimates (Hänfling et al., 2016; Lawson Handley et al., 2019; Sard et al., 2019).

Current uncertainties regarding the quantitative power of eDNA metabarcoding ultimately originate from our lack of knowledge on the origin and fate of eDNA in aquatic systems (Klymus et al., 2015; Lacoursière-Roussel et al., 2016; Sassoubre et al., 2016). Age, physiology, life history and metabolic rate all play a role in the amount of eDNA released (eDNA shedding rate) from organisms into their surroundings (Barnes et al., 2014; Goldberg et al., 2016; Ruppert et al., 2019). Physical, chemical and biological forces such as dilution, sedimentation and resuspension, hydrolysis, oxidation and microbial activity, can all influence eDNA persistence and dynamics within aquatic habitats (Turner et al., 2015). In addition, the degradation of genetic material is also promoted by high temperature and acidity (Seymour et al., 2018; Ruppert et al., 2019). In our study system, the fish age distribution was relatively narrow, therefore, reducing the effect of different eDNA shedding rates from distinct life stages and age classes. Moreover, the ponds surveyed were similar in terms of high fish density and species composition and were also exposed to stable environmental conditions that positively influenced the reproducible and reliable quantitative characterisation of the fish communities investigated.

A lack of robust sampling and metabarcoding protocols may also contribute to a distortion of the observed diversity patterns. Insufficient sampling effort, inhibition, primer biases, sequencing artefacts, database inaccuracy and contamination are the main methodological sources of bias (McKee et al., 2015; Grey et al., 2018; Collins et al., 2019; Wood et al., 2019). In the present study, the quantitative fish assessment of the two ponds surveyed demonstrates the accuracy of optimized eDNA metabarcoding protocols to reflect species biomass and abundance. In recent years, sampling, laboratory, and bioinformatics workflows have been progressively refined for the characterization of fish communities within UK freshwater ecosystems (Hänfling et al., 2016; Li et al., 2018; Lawson Handley et al., 2019; Li et al., 2019a). Here, we have demonstrated that optimised sampling strategy, enhanced extraction protocol with an additional inhibitor removal step (Sellers et al., 2018), replication during PCRs and development of a custom-curated database with *de novo* sequences, strengthened the probability of detection, reduce taxonomic assignment bias, and overall provided reliable quantitative data of fish biomass and number of individuals.

### Impact of DNA capture and preservation methods

In our study the correlations between sequence read counts and species abundance/biomass were consistently high for all filtration treatments with average correlation coefficients of 0.93 for Sterivex filters and 0.84 for MCE filters (Fig. 3). The variation of correlations observed between filter types may be explained from differences of read counts assigned to species as a result of different water volumes filtered between Sterivex and MCE filters (Fig. S1). However, for equal amounts of water filtered and high DNA concentrations, open filter membranes usually capture a higher amount of DNA compared to enclosed filters possibly due to the tendency of enclosed filters to clog more easily (Li et al., 2018; Takahashi et al., 2020). Quantitative differences between filter types may also vary with the target species as observed in this study (Takahashi et al., 2020). In fact, while we observed a general trend of higher species read counts in MCE filters, we also observed the opposite trend for *C. carpio* which showed lower reads in MCE filter replicates compared to Sterivex filters (Fig. S1).

The higher species richness found in Sterivex filters preserved with RNAlater and open filter membrane replicate 1 resulted from the detection of only one low-abundant taxa within the pond (*S. erythropthalmus*; Fig. 1). We therefore consider this result a stochastic effect between filter replicates or storage methods.

Overall, we found that both filter types showed a good representation of fish diversity and community composition and, consequently, we suggest that they can be used interchangeably depending on time, resources and location of the study. Sterivex filters, for instance, are effective for field processing of water samples, facilitating collection in remote locations. After sample collection, Sterivex are immediately filtered on-site (using peristaltic pumps or sterile syringes) and the risk of contamination is reduced because of the lack of filter handling (Spens et al., 2017; Li et al., 2018). In the present study, there was no evidence of higher contamination in open filter membranes compared to Sterivex filters indicating that preventing on-site and in-lab contaminations is sufficient to minimise/avoid DNA contaminations regardless of the filter types choice. The use of Sterivex filters, or enclosed filters in general, is however more amenable to large scale monitoring programs for environmental managers or citizen science projects (Biggs et al., 2015; Buxton et al., 2018; Larson et al., 2020). Nevertheless, Sterivex filters are currently almost 15 times more expensive than open filters, DNA extraction is more time-consuming, and, when syringes are used for filtration, the Sterivex method requires a large amount of disposable plastic consumables. The use of prepacked sterile syringes is nonetheless preferred over pumps suction (vacuum or peristaltic) to reduce filtration time (this paper; Li et al., 2018).

## Conclusion

This study underpins valuable considerations for the quantitative estimates of eDNA metabarcoding data. We demonstrated that eDNA metabarcoding data correlate with actual abundance and biomass of fish communities within small freshwater systems with high fish density.

Established methods (i.e. hydroacustic, electrofishing, gillnetting) for obtaining quantitative estimates of fish abundance are resource intensive and may not be suitable for all water bodies and species (Winfield et al., 2009). Furthermore, quantitative interpretation of data is often complex (hydroacustic) or relies on large sampling effort (netting/electrofishing) (Winfield et al., 2009), hence becoming costly in terms of financial, human resources and habitat disturbance or species mortality. More importantly, these methods can be prone to errors as they are not exhaustive sampling methods and, therefore, can only provide approximation of species abundance.

eDNA metabarcoding is arguably a more flexible tool, adaptable to all aquatic environments and fish species, is non-lethal, and the sources of errors can be minimized through a careful optimization of field and laboratory protocols.

Monitoring trends in population size and community structure is paramount to the assessment of species health and viability, and the outputs are required to undertake management actions and to guide conservation decisions (Kull et al., 2008). Implementation of eDNA metabarcoding will drive a step-change towards non-invasive monitoring strategies for next-generation ecosystems surveillance. eDNA metabarcoding, as a non-invasive, fast, universally applicable approach, is nowadays claiming the attention of researchers, stakeholders and governmental agencies. Therefore, exploring, evaluating and finally establishing the quantitative value of such a broadly-used tool for diversity monitoring is essential.

## Acknowledgements

This work was funded by the UK Environment Agency (collaborative agreement 171024). We would like to express our thanks to the EA national and local staff working at the site during the eradication program and the fishery farm owner for providing us the opportunity of using the site for this research study. We are also very grateful to Peter Shum, Jairo Arroyave and Stephanie McLean for the constructive feedback on the manuscript before its initial submission.

