## Supplementary figure 1 and table 1 for "Read counts from environmental DNA (eDNA) metabarcoding reflect fish abundance and biomass in drained ponds"

**Table S1.** List of fish species and their abundance/biomass present in New Lake (NL) and Middle Lake (ML) at the fishery farm.

| Pond | Common name | Scientific name | Number of individuals | Total biomass (kg) |
|---|---|---|---|---|
| NL | Common carp | *Cyprinus carpio* | 293 | 852.8 |
| NL | Hybrid (Common carp x Crucian carp) | *Cyprinus x Crucian* | 17 | 16 |
| NL/ML | Bream | *Abramis brama* | 382 | 240.6 |
| NL/ML | Barbel | *Barbus barbus* | 3 | 3.1 |
| NL/ML | Crucian carp | *Carassius carassius* | 378 | 123.7 |
| NL/ML | Hybrid (Common carp x Crucian carp) | *Cyprinus x Crucian* | 7 | 2.9 |
| NL/ML | Perch | *Perca fluviatilis* | 23 | 10.57 |
| NL/ML | Hybrid (Roach x Bream) | *Rutilus x Abramis* | 60 | 31.4 |
| NL/ML | Roach | *Rutilus rutilus* | 184 | 46.9 |
| NL/ML | Rudd | *Scardinius erythrophthalmus* | 7 | 1.5 |

| | | | | |
|---|---|---|---|---|
| NL/ML | Chub | *Squalius cephalus* | 1 | 0.7 |
| NL/ML | Tench | *Tinca tinca* | 203 | 173.5 |
| NL | Crucian carp | *Carassius carassius* | 1 | 0.2 |
| NL | Grass carp | *Ctenopharyngodon idella* | 4 | 17.9 |
| NL | Common carp | *Cyprinus carpio* | 190 | 862.4 |
| NL | Hybrid<br>(Common carp x Crucian carp) | *Cyprinus x Crucian* | 4 | 1.8 |
| NL | Perch | *Perca fluviatilis* | 5 | 1.2 |
| NL | Bream | *Abramis brama* | 151 | 42.7 |
| NL | Sterlet | *Acipenser spp.* | 1 | 1 |
| NL | Hybrid (Roach x Bream) | *Rutilus x Abramis* | 5 | 1.8 |
| NL | Roach | *Rutilus rutilus* | 58 | 10.7 |
| NL | Wels catfish | *Silurus glanis* | 23 | 251.95 |

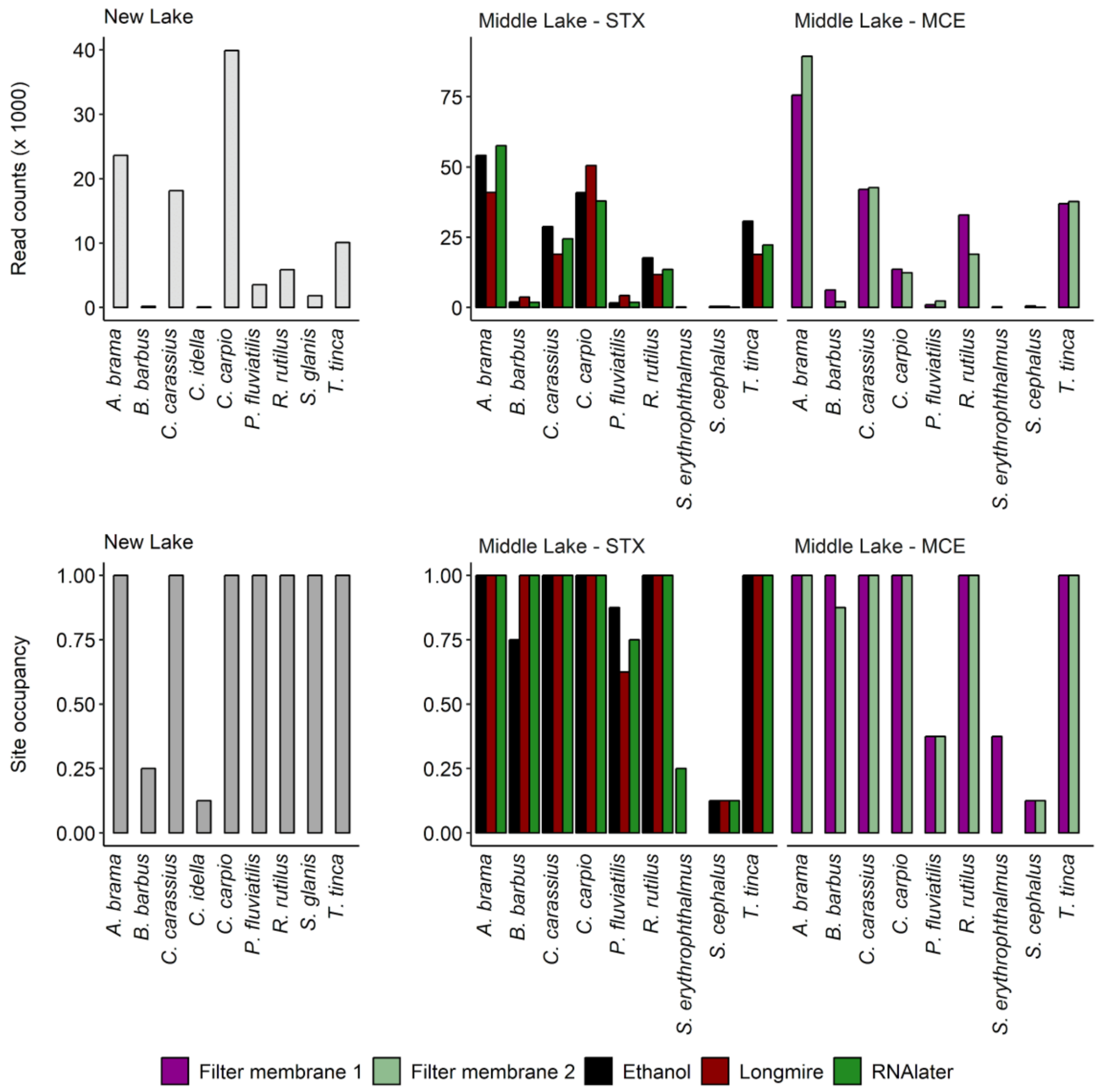


**Figure S1.** Fish species read counts (top) and site occupancy (bottom) barplots for New Lake, Middle Lake Sterivex sampling (Middle Lake-STX) and Middle Lake open filters sampling (Middle Lake-MCE). *Note* MIX samples were not included in site occupancy calculations.
